# A probable genetic origin for pit defects on the molars of *Paranthropus robustus*

**DOI:** 10.1101/400671

**Authors:** Ian Towle, Joel D. Irish

**Affiliations:** Research Centre in Evolutionary Anthropology and Palaeoecology, School of Natural Sciences and Psychology, John Moores University, Liverpool, United Kingdom, L3 3AF

**Keywords:** enamel hypoplasia, primary teeth, dental defects, amelogenesis imperfecta

## Abstract

We report the frequencies of linear enamel hypoplasia (LEH) and, specifically, pitting enamel hypoplasia (PEH) defects in the teeth of *Paranthropus robustus*, for comparison with four other South African hominin species and three extant nonhuman primate species. Unlike LEH, the lesser known PEH is characterized by multiple circular depression defects across a tooth crown and is often difficult to interpret in terms of developmental timing and etiology. Teeth in all samples were examined macroscopically with type, position and number of defects recorded. Frequencies of teeth with LEH vary among hominin species, but the differences in PEH are considerable. That is, *P. robustus* has much higher rates of pitting defects, with 47% of deciduous teeth and 14% of permanent teeth affected, relative to 6.7% and 4.3%, respectively, for all other hominin teeth combined; none of the extant primate samples evidence comparable rates. The defects on *P. robustus* molars are unlike those in other species, with entire crowns often covered in small circular depressions. The PEH is most consistent with modern human examples of amelogenesis imperfecta. Additionally, the defects are: 1) not found on anterior teeth, 2) uniform in shape and size, and 3) similar in appearance/severity on all molars. A possible reason for this form of PEH is as a side effect of selection on another phenotype that shares the same coding gene(s), i.e., a genetic origin. Recent research on the ENAM gene provides one such possibility. *Paranthropus* likely underwent rapid evolution in the ENAM loci, with changes in this gene contributing to larger posterior teeth and thicker enamel. This same gene is associated with amelogenesis imperfecta; therefore, pleiotropy effects, relating to high selection on this gene during *Paranthropus* evolution, could have yielded this unique condition.

## Introduction

Enamel hypoplasia occurs during the secretory stage of formation, whereas other enamel defects form during the maturation stage, e.g., hypocalcification and dental fluorosis (Ten Cate, 1994; Guatelli-Steinberg, 2015; Xing et al., 2015). Defects take a variety of forms, most of which have been found in fossil hominins (e.g., Tobias, 1967; Goodman et al., 1987; Moggi-Cecchi, 2000; Lukacs et al., 2001; Guatelli-Steinberg et al., 2004; Xing et al., 2015). A range of disturbances can create similar defects, often making a diagnosis of particular etiologies difficult. Nonetheless, enamel hypoplasia may be able to provide some insight into diet, genetic conditions, environment factors and health of individuals and populations (e.g., Cunha, 2004; Ogden et al., 2007; Schuurs, 2012; Guatelli-Steinberg et al., 2014).

Enamel hypoplasia is often split into three broad categories: linear (LEH), pit (PEH) and plane-form (Pindborg, 1970; Seow, 1990). Defects can look remarkably different, but ultimately all are associated with a reduction of enamel matrix from disruption in ameloblast production (Eversole, 1984). It is not always easy to assign defects to these categories (e.g., Ogden, 2007), but doing so is often justified because they can have specific etiologies. In particular, genetic conditions, injuries to the tooth during formation, and certain diseases can cause characteristic hypoplasia defects (Cook, 1980; Goodman & Rose, 1991; Skinner & Newell, 2003; Weerheijm, 2003; Crawford et al., 2007; Ogden et al., 2008).

Methods for recording enamel hypoplasia vary among studies. Most researchers record and compare LEH frequencies (e.g., Guatelli-Steinberg, 2004; Miszkiewicz, 2015; Smith et al., 2016). Others include all defects (e.g., Goodman et al., 1980, 1984; Goodman & Armelagos, 1985; Ogilvie et al., 1989). Some studies only record defects on certain teeth, with posterior and deciduous teeth excluded (e.g., Infante & Gillespie, 1974; Lovell & Whyte, 1999). Additionally, it is not always clear if the pitting hypoplasia mentioned only refers to defects found as part of LEH bands (e.g., Mellanby, 1929; Sognnaes, 1956; Goodman et al., 1980, 1984; Goodman & Armelagos, 1985; Hillson, 1992).

Pitting enamel hypoplasia may take a variety of forms, ranging from small circular pinpricks to vast irregular depressions (Skinner, 1996; Hillson & Bond, 1997; Witzel et al., 2006; Ogden, 2007). Pits also vary in distribution across a tooth crown, with some forming rows around the circumference, usually associated with shallow defects, and others more randomly scattered (Goodman & Rose, 1990). PEH can also be associated with plane-form hypoplasia (e.g., Ogden et al., 2007; Lauc et al., 2015), though commonly it is the only defect observed.

The location of PEH does not necessarily give insight into the age of the individual when the defect formed. The reason is that pit depth is related to its position on the plane of the brown striae of Retzius on which enamel matrix formation ceased. Deep pits may, therefore, represent a disturbance much earlier than their crown position suggests. A further issue in studying PEH is that it is not yet clear why it forms instead of other hypoplasia types, particularly LEH. However, the tooth involved, position on the crown, and cause of the disruption are all important factors. Typically, only the occlusal type of perikymata is affected. Molars have significantly more of their crown surface covered with this type, which may explain why they often have more PEH (Hillson & Bond, 1997; Hillson, 2014). However, it has also been suggested that because it is uncommon for an individual to have both LEH and PEH, these different hypoplasia types may have different etiologies (Lovell & Whyte, 1999).

Each pit corresponds to the ceasing of ameloblast activity at a particular point in enamel formation. However, it is not clear why only certain ameloblasts are affected along the plane of a brown stria of Retzius during formation (Witzel et al., 2006; Ogden et al., 2007). In some cases, only a few ameloblasts stop forming enamel matrix, leading to small pits. With large pits, hundreds of these enamel-forming cells may cease production (Guatelli-Steinberg, 2015). In other forms of systemic enamel hypoplasia, such as LEH and plane form, all ameloblast activity is affected. In most instances of PEH the enamel between pits appears normal. Exposed Tomes’ process pits can often be observed within pits, showing a sharp end to the ameloblasts; however, some examples show continued deposition of irregular enamel (Hillson & Bond, 1997; Witzel et al., 2006).

In most cases of PEH described in the literature, particularly in archaeological examples, researchers have not been able to specify a cause; instead, some form of non-specific stress was suggested. Nevertheless, PEH has been associated with a number of specific disturbances in modern clinical studies, including: hypocalcaemia, premature birth, low birth weight, hypoparathyroidism, neonatal tetany, maternal diabetes mellitus, kernicterus, vitamin D deficiency, congenital syphilis, amelogenesis imperfecta, and nutritional deficiency (Eliot et al., 1934; Grahnen & Selander, 1954; Croft et al., 1965; Purvis et al., 1973; Stimmler et al., 1973; Pisanty et al., 1977; Nikiforuk & Fraser, 1979, 1981; Seow et al., 1984; Wright et al., 1993; Aine et al., 2000; Pinhasi et al., 2006; Gaul et al., 2015; Radu & Soficaru, 2016).

Differences in PEH frequencies among fossil hominins and extant primates have rarely been explored. Pitting enamel hypoplasia has, however, been found on the teeth of various hominin specimens (e.g., Tobias, 1967; Ogilvie et al., 1989; Xing et al., 2015; Zanolli et al., 2016). Some studies have noted the presence of PEH in *P. robustus* teeth (Robinson, 1956; White, 1978; Moggi-Cecchi, 2000; Moggi-Cecchi et al., 2010). These South African hominin studies have been in the context of hypoplasia rates as a whole, and to date a cause for these defects has not been explored. In the present study, PEH frequencies and appearance in *P. robustus* will be compared with those in other hominins and extant primates; a differential diagnosis to explain these unusual defects will follow.

## Materials and Methods

Of 431 *P. robustus* teeth, 127 (29.47%) could not be recorded for hypoplasia due to crown damage or post-mortem discoloration, leaving 304 teeth for the study. The comparative material includes specimens assigned to Early *Homo, Australopithecus sediba, Homo naledi, A. africanus*, gorillas, chimpanzees and baboons (Table 1). The South African fossil hominin samples are curated at the University of the Witwatersrand and The Ditsong National Museum of Natural History. The extant primate samples are curated at the Powell-Cotton Museum. All of the latter individuals were killed in their natural habitat and comprise western lowland gorillas (*Gorilla gorilla gorilla*), common chimpanzees (*Pan troglodytes*) and olive baboons (*Papio anubis*) (Dean & Jones, 1992; Guatelli-Steinberg & Skinner, 2000; Lukacs, 2001).

**Table 1.**
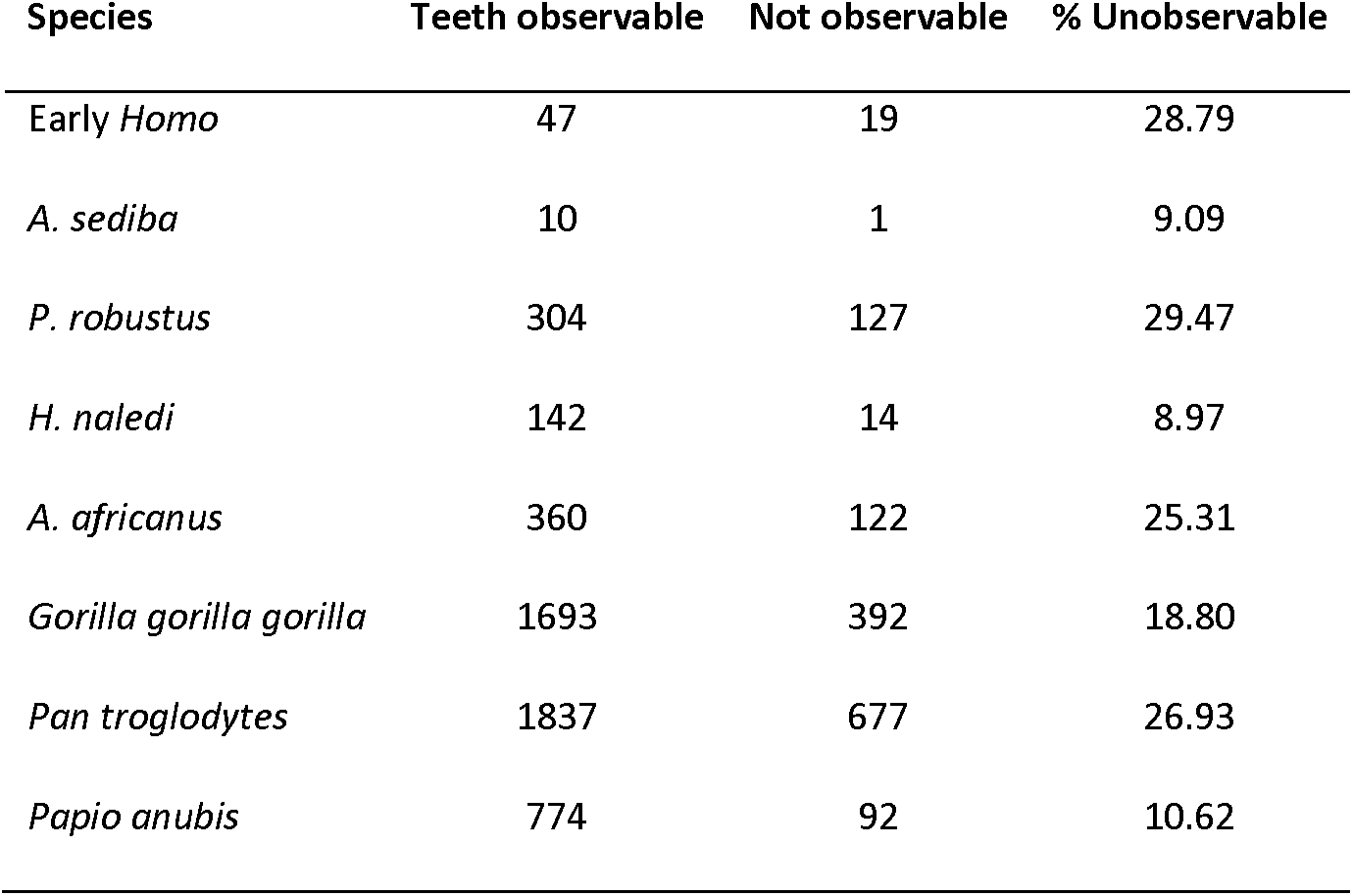
Number of observable teeth for each species.

Teeth were observed macroscopically, with a 10x hand lens used to clarify defect types. The presence and position of linear, localized, pitting and plane-form hypoplasia were recorded for each tooth, along with defect size and shape. All teeth were viewed under an incandescent lamp and slowly rotated so light hit at a variety of angles to discern even the smallest of defects. The presence of PEH was recorded, and notes on position described. If pits were part of an LEH defect, the position of the LEH was recorded and pitting noted; however, this form of pitting is not included in the PEH analysis. To compare certain defect types, as well as between species, a chi-square test of homogeneity with significance set at the 0.05 alpha level was used.

In some cases, micro-CT scans of *P. robustus* molars were viewed and defect size and number recorded. In particular, CT scans of six teeth with defined defects were viewed to take precise measurements of pit diameter and get a better understanding of defect depth and uniformity. The most defined pits that show up on the CT scanned 3D model were recorded after corroboration with photos of the specimen. Between 15 and 20 defects were measured for each tooth, with mean and range recorded. The scans were completed by the Department of Human Evolution, Max Planck Institute for Evolutionary Anthropology with a BIR ACTIS 225/300 (kV, 100 mA, 0.25 brass filter) or a SkyScan 1172 (100 kV, 94 mA, 2.0 mm aluminium and copper filter) microtomographic scanner. The isometric voxel sizes resulting from these scans range between 15 and 50 micrometres (mm) (Skinner, personal communication, 2018).

Data are presented by tooth count, where the number of hypoplastic teeth is displayed as a percentage of the total number of observable teeth. Because the hominin material is fragmentary, this approach yields maximum sample size for comparison; additionally, Lovell and Whyte (1999) note that displaying on a ‘per-tooth’ rather than ‘per-individual’ basis also permits a broader comparison of subsamples of different tooth groups among populations. Antimeres are treated as separate data points in the overall hypoplasia frequencies. This strategy increases sample sizes and allows maximum recovery of information. Two additional points justify including antimeres: 1) some defects may be displayed on one side and not the other, in particular, localized defects, and 2) the nature of the fossil record means that, in some cases, correctly assigning antimeres is difficult.

Instead of rejecting teeth worn past a certain point, all crowns that are unbroken due to post-mortem damage were included. This approach allows inclusion of teeth that may have had enamel defects worn away during life. However, excluding worn teeth also leads to bias, since an entire sample would consist of individuals who died young, i.e., potentially more ill, on average, during dental development than those who lived to old age. Additionally, many defects remain visible even when the tooth has significant wear, particularly PEH defects in *P. robustus*.

## Results

The *P. robustus* sample exhibits an extremely high rate of PEH, far higher than any extant primate or fossil hominin sample studied (Tables 2 and 3). There is a statistically significant difference in PEH in permanent teeth between *P. robustus* and the other hominins and primates (e.g., *P. robustus* vs. *A. africanus* X^2^= 14.823, 1 df, p= 0.0001). The case is similar for deciduous teeth (*P. robustus* vs. *A. africanus* X^2^= 5.824, 1 df, p= 0.0158).

**Table 2.** Per tooth frequencies (%) of Pitting Enamel Hypoplasia (PEH), Linear Enamel Hypoplasia (LEH), and localized hypoplasia for permanent teeth of each species.

| Species | PEH (# teeth) | LEH (# teeth) | Localized (# teeth) |
| --- | --- | --- | --- |
| *Pan troglodytes* | 0.65 (12/1837) | 8.06 (148/1837) | 0.98 (18/1837) |
| *Gorilla gorilla gorilla* | 2.89 (49/1693) | 4.25 (72/1693) | 0.95 (16/1693) |
| *Papio anubis* | 0.00 (0/774) | 2.07 (16/774) | 1.68 (13/774) |
| *H. naledi* | 0.70 (1/142) | 14.79 (21/142) | 0.70 (1/142) |
| *A. africanus* | 5.03 (18/358) | 15.08 (54/358) | 0.28 (1/358) |
| *P. robustus* | 14.75 (41/278) | 11.51 (32/278) | 1.08 (3/278) |
| Early *Homo* | 0.00 (0/47) | 8.51 (4/47) | 2.13 (1/47) |
| *A. sediba* | 0.00 (0/10) | 0.00 (0/10) | 0.00 (0/10) |

**Table 3.** Per tooth frequencies (%) of Pitting Enamel Hypoplasia (PEH) and localized hypoplasia for deciduous teeth of each species.

| Species | PEH (# teeth) | Localized (# teeth) |
| --- | --- | --- |
| *Pan troglodytes* | 4.23 (25/591) | 5.08 (30/591) |
| *Gorilla gorilla gorilla* | 1.39 (6/433) | 12.93 (56/433) |
| *Papio anubis* | 0.00 (0/107) | 3.74 (4/107) |
| *H. naledi* | 0.00 (0/16) | 0.00 (0/16) |
| *A. africanus* | 5.00 (2/19) | 0.00 (0/19) |
| *P. robustus* | 41.30 (19/46) | 0.00 (0/46) |
| Early *Homo* | 14.29 (2/14) | 0.00 (0/14) |

The first, second and third permanent molars of *P. robustus* are similarly affected, with over 20% displaying PEH defects. Therefore, the high PEH rates on permanent teeth of *P. robustus* are related predominately to molars, although premolars also exhibit high rates relative to the other hominin species (Table 4). For the permanent dentition as a whole, *P. robustus* has more teeth with PEH than LEH.

**Table 4.** Per tooth frequencies (%) for Pitting Enamel Hypoplasia (PEH) and Linear Enamel Hypoplasia (LEH) in the three largest hominin samples, for different tooth groups of permanent teeth.

| Permanent teeth | *P. robustus* |  | *A. africanus* |  | *H. naledi* |  |
| --- | --- | --- | --- | --- | --- | --- |
|  | PEH | LEH | PEH | LEH | PEH | LEH |
| Anterior teeth | 1.75 (1/57) | 22.81 (13/57) | 2.11 (2/95) | 40.00 (38/95) | 2.04 (1/49) | 24.49 (12/49) |
| Premolars | 7.04 (5/71) | 9.86 (7/71) | 0.00 (0/90) | 6.67 (6/90) | 0.00 (0/39) | 23.08 (9/39) |
| First molars | 21.43 (12/56) | 5.36 (3/56) | 6.90 (4/58) | 5.17 (3/58) | 0.00 (0/25) | 0.00 (0/25) |
| Second molars | 21.57 (11/51) | 5.88 (3/51) | 9.09 (6/66) | 6.06 (4/66) | 0.00 (0/19) | 0.00 (0/19) |
| Third molars | 25.58 (11/43) | 11.63 (5/43) | 10.20 (5/49) | 4.08 (2/49) | 0.00 (0/10) | 0.00 (0/10) |

In the deciduous teeth of *P. robustus*, PEH defects also occur primarily on molars (Table 5). The first and second deciduous molars, in both jaws, are similarly affected, with PEH present on 54% and 52% of these teeth, respectively; indeed, the crown is often completely covered, to resemble dimples on a golf ball (Figure 1). On both permanent and deciduous teeth of *P. robustus*, severe PEH often covers large areas of the crown, and characteristically comprises numerous uniform small depressions. Pits are typically more defined toward the occlusal surface. Defects vary little in size or shape, and hundreds of separate pits are often visible across the crown, with nearly identical defects evident in both permanent and deciduous molars (Figure 2; Table 6).

**Table 5.** Percentage of deciduous *P. robustus* teeth with Pitting Enamel Hypoplasia (PEH).

| Deciduous Teeth | PEH % | Total teeth | Teeth with hypoplasia |
| --- | --- | --- | --- |
| All teeth | 41.30 | 46 | 19 |
| Anterior teeth | 8.33 | 12 | 1 |
| First molars | 53.85 | 13 | 7 |
| Second molars | 52.38 | 21 | 11 |

**Table 6.** Number and size of *P. robustus* pitting defects. Pit size is in millimeters. L: left; R: right; d: deciduous; M1: first molar; M2: second molar. A total of 15-20 of the most defined pits was analyzed for each specimen. All mandibular teeth except SK 89.

| Specimen | Number of pits | Minimum pit size | Maximum pit size | Average |
| --- | --- | --- | --- | --- |
| SK 61: RM1 | 50+ | 0.12 | 0.36 | 0.23 |
| SK 61: RdM2 | 80+ | 0.11 | 0.28 | 0.17 |
| SK 61: LdM2 | 80+ | 0.12 | 0.26 | 0.18 |
| SK 63: RdM2 | 100+ | 0.11 | 0.42 | 0.23 |
| SK 64: RdM2 | 300+ | 0.12 | 0.21 | 0.16 |
| SK 89: LM1 | 100+ | 0.12 | 0.26 | 0.16 |

**Figure 1.**
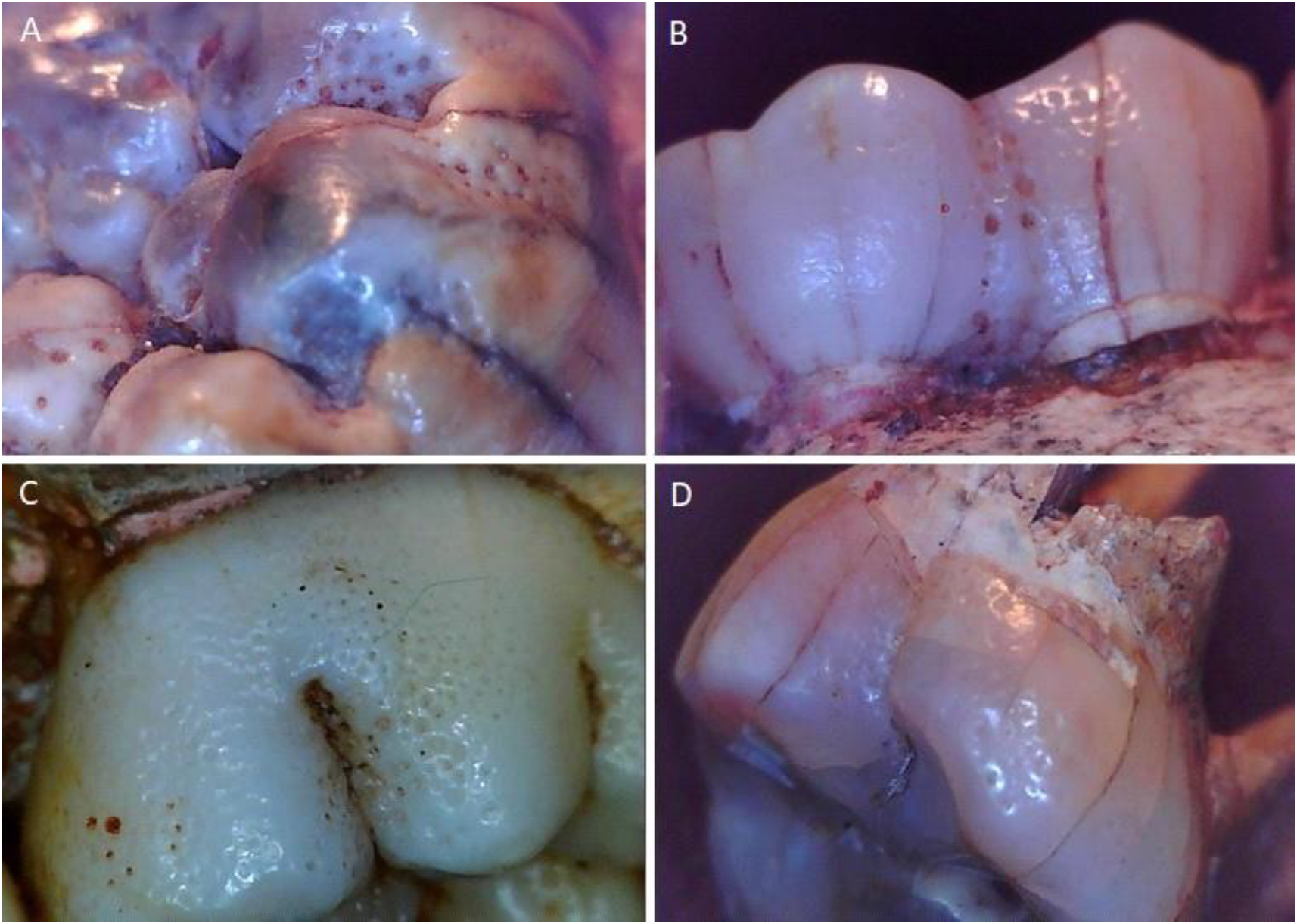
Uniform circular pitting enamel hypoplasia on four *P. robustus* teeth. A) SK 61; B) SK 63; C) SK 64; D) SK 90.

**Figure 2.**
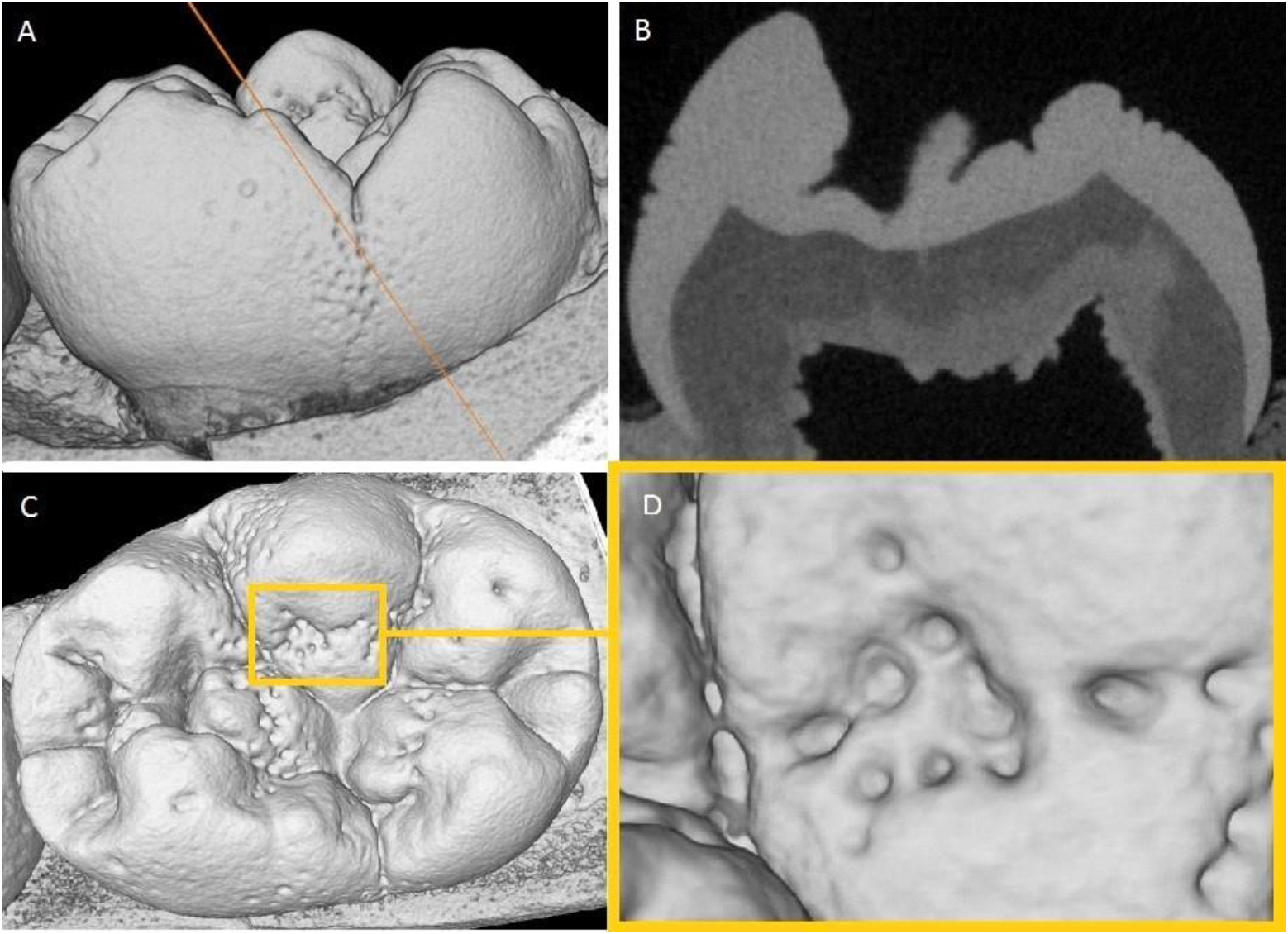
Micro-CT scan slices of SK 64 (*P. robustus*), right mandibular first molar. A) Overview of the lingual surface with the orange line the position of the slice in B; B) Slice showing pitting on both the lingual and buccal surfaces. C) Occlusal view; D) Close up of pitting on the occlusal surface.

In deciduous molar antimeres with PEH, both tend to have mirror-image defects. For example, the right and left second deciduous molars of SK 61 not only have a similar distribution of uniform pits, but areas where the most defined defects are present is identical. These pits were clearly not made post-mortem, and the surrounding enamel does not seem to have been reduced, i.e., it appears to be of normal density (Figure 2). The rate of PEH is similar between *P. robustus* sites. At Kromdraii 16.67% of permanent teeth have PEH, while at Swartkrans this figure is 14.34%. Anterior teeth associated with affected permanent molars do not show an increase in LEH defects, as only 16.67% of these teeth are affected.

## Discussion

Few studies have reported on different types of hypoplasia, and, in particular, it is uncommon to report PEH frequencies. Lovell and Whyte (1999) studied a human sample from Ancient Mendes, Egypt, finding that linear defects were over three times more common than pitting defects. However, their permanent teeth samples only consisted of anterior teeth. Similarly, Goodman et al. (1987) compared frequencies of pitting between deciduous and permanent anterior teeth, finding higher frequencies in the latter. In a modern sample, Pedersen (1944) found 14% of two to four-year-old children had enamel hypoplasia in their teeth. Typically, in human samples, the frequency of hypoplasia in deciduous teeth is ≤ 5% (e.g., Lovell & Whyte, 1999; Robles et al., 2013). However, studies focusing on humans severely affected by disease, famine, or malnutrition show much higher rates, with 18% to 62% of teeth affected (Enwonwu, 1973; Infante & Gillespie, 1974; Seow, 1990). Ogilvie et al. (1989) found a high rate in Neanderthals. Half of those defects on the posterior teeth were of the PEH variety. Deciduous teeth also showed pitting, but at a rate of just 3.9%. The authors note that later developing Neanderthal teeth seem more affected, with an increasing number of defects from the first to third molars. This low frequency in deciduous teeth and permanent first molars stands in stark contrast to that of *P. robustus,* with deciduous molars most affected.

Due to the lack of known frequencies in other samples, it is not known how common PEH is in hominins overall. That said, it is clear from the present results, as well as in studies of modern humans, that the PEH rate in *P. robustus* is remarkably high (Goodman et al., 1987; Ogilvie et al., 1989; Seow et al., 1992). Such defects are also evident on *P. robustus* posterior teeth not included in this study, from Drimolen and Cooper’s, as well as similar PEH in *P. boisei* (Tobias, 1967; de Ruiter et al., 2009; Moggi-Cecchi et al., 2010). The frequency of these defects across sites suggests *P. robustus,* and perhaps the greater *Paranthropus* genus, shared this tendency for ‘golf ball-like’ enamel defects.

Extant ape species studied to date do not have high PEH rates, and the appearance of defects varies from those in *P. robustus*. Additionally, the PEH on *P. robustus* molars does not resemble hypoplastic defects in many modern clinical studies of deciduous teeth, including those caused by premature birth, low birth weight, vitamin D deficiency, tuberous sclerosis, congenital syphilis, pseudohypoparathyrgidism and epidermolysis bullosa (Croft et al., 1965; Purvis et al., 1973; Stimmler et al., 1973; Nikiforuk & Fraser, 1979, 1981; Seow et al., 1984; Wright et al., 1993; Aine et al., 2000; Pinhasi et al., 2006; Gaul et al., 2015; Radu & Soficaru, 2016). Most of these conditions are associated with pitting that is irregular in both shape and distribution. They also tend to affect all teeth, not just molars, and are associated with other types of dental defects.

Pitting defects have also been associated with deficiencies or surpluses of certain compounds. For example, dental fluorosis can create pit like features on tooth crowns (Fejerskov et al., 1990); they can manifest in several ways but mostly as white opaque lines that, in extreme cases, can lend a chalky white appearance to the entire crown (Fejerskov et al., 1990; Xing et al., 2015). If the fluorosis is severe, the outer enamel may eventually fracture to leave cracks or pits— though the latter have edges that appear jagged or broken under magnification (Thylstrup & Fejerskov, 1978). Another effect of dental fluorosis, particularly in severe cases, is discoloration. The *P. robustus* teeth do not show any of these features, so fluorosis is unlikely. Other compounds, e.g., mercury, can create enamel defects during dental development (Ogden, 2007; Ioannou et al., 2015; Radu & Soficaru, 2016). Similarly, a common cause of prenatal enamel hypoplasia has been linked to calcium deficiency in the mother, from malnutrition or malabsorption (Lovell & Whyte, 1999). Postnatal defects have also been linked to hypocalcaemia, from insufficient calcium consumption or malabsorption. The defects in *P. robustus* do appear consistent with such an etiology, so perhaps they result from a specific environmental or dietary component, or absence thereof. That said, the lack of affected anterior teeth is not supportive of this scenario.

Ogden et al. (2007) describe a human sample with high hypoplasia rates in deciduous and permanent dentitions from 16th–18th Century London. Superficially, some defects look similar to the pitting on *P. robustus* molars. Yet, the pitting is less uniform, with evidence of both plane-form and pitted defects. Additionally, the cusp morphology in the London sample is affected in many teeth, which does not appear to be the case for *P. robustus*. The example in Hillson (2014) of PEH in a modern deciduous first molar and plane form defects on the second deciduous molar and canine, is suggested as a case of ‘neonatal dental hypoplasia’, in which a stress at birth is the cause. Thus, a potential hypothesis is that the *P. robustus* defects form at or around birth, perhaps from an environmental or nutritional disturbance. However, two issues counter this hypothesis: 1) the lack of associated hypoplasia on other deciduous teeth, and 2) pitting is equally present on all permanent molars.

A neonatal line can be found in a variety of populations, affecting those teeth forming at birth; in humans, this means the deciduous dentition and first permanent molars. It is thought to result from either a decrease in plasma calcium after birth (Norén et al., 1984; Smith & Avishai, 2005) or disturbances from the birth process (Whittaker & Richards, 1978; Guatelli-Steinberg, 2015). Although not explored in many fossil hominin samples, if neonatal lines are found in the deciduous and permanent first molars of *P. robustus* with pitting, inferences into timing may be permitted. Unfortunately, at present, the only *P. robustus* tooth for which these data are available is not pitted (Smith et al., 2015). If the defects are not genetic in origin, and at least some PEH defects are formed in utero, then it likely relates to nutritional factors in the mother (Seow, 1990; Lukacs, 1992; Lovell & Whyte, 1999). Clearly, however, PEH defects on second and third permanent molars indicate a postnatal etiology.

Perhaps the thick enamel and faster development of *Paranthropus* molars compared to other hominin species (Lacruz et al., 2008) meant that they were more predisposed to developmental defects. For example, the shape of the crown and the structure of the enamel may have meant that even a slight disturbance during development resulted in severe defects. A single brown stria of Retzius affected by pockets of ameloblasts ceasing enamel matrix formation could potentially yield surprisingly large areas of PEH (Hillson and Bond, 1997; Witzel et al., 2006). Therefore, PEH in *P. robustus*, in which the entire crown is covered in small uniform depressions, could potentially have been caused by a relatively brief disturbance. Brich & Dean (2013) found that in modern humans: 1) second deciduous molars form enamel at a slower rate than other deciduous teeth, and 2) the cusps of deciduous second molars have thicker enamel and take proportionally longer to form. Disturbances occurring whilst molar cusps are still forming can lead to pitting defects over large areas of the crown (Hillson, 2014). Therefore, it is a possibility that PEH defects on *P. robustus* deciduous molars may relate to the thick enamel and developmental timing of these teeth. This possibility would explain why defects are much less common on the anterior teeth. However, a disturbance event is still necessary to cause the defects, so the uniform appearance and common occurrence makes this hypothesis unlikely.

The pitting in *P. robustus* resemble certain modern cases of amelogenesis imperfecta. This set of genetic disorders affects one in every 700 to 14,000 modern humans (Sundell & Koch, 1984; Crawford et al., 2007). Defects usually include scattered pits and plane form hypoplasia but abnormal colouration, thickness and density of the enamel is also common (Wright, 1985; Aldred et al., 2003; Chamarthi et al., 2012; Schuurs, 2012; Towle et al., 2017). A specific type called hypoplastic amelogenesis imperfecta (Wright, 1993; Mehta et al., 2013) most closely emulates the defects in *P. robustus*, in that it commonly does not show coloration, thickness and density abnormalities of the enamel (Witkop & Sauk, 1976; Witkop, 1988; Seow, 1992). Indeed, the pitting defects look very similar to some examples of this type of amelogenesis imperfecta (e.g., Figure 3 of Rushton, 1964; Ozdemir et al., 2005). A recent study highlighted a particular genetic mutation that causes defects to be present on posterior, but not anterior primary teeth (Kim et al., 2016). Therefore, it may be possible that the PEH on *P. robustus* molars is genetic in origin. The fact that both deciduous and permanent molars are similarly affected and that all defects are essentially identical in appearance supports this possibility.

Of course, it would be important to understand the reason(s) for such a high rate of ‘genetically-related’ enamel defects in *P. robustus* from an evolutionary perspective. Enamel hypoplasia predisposes teeth to caries and extreme wear. Yet, specimens assigned to *P. robustus* with PEH from different sites, along with potentially similar pitting in east African *Paranthropus* specimens, suggest this type of defect was common for an extended period of time.

Recent advances in genetic analyses of dental development may suggest a likely scenario. Rapid evolution of a particular feature can lead to loss of genomic stability in particular genes, or create pleiotropy effects on other characteristics (Pavlicev & Cheverud, 2015; Fiddes et al., 2018; Hlusko et al., 2018). For example, Hlusko et al. (2018) found the EDAR gene to be under intense selection in certain populations during the last ice age, relating to transmission of vitamin D and fatty acids; the same populations show specific dental characteristics associated with changes in this gene. Perhaps the *Paranthropus* PEH defects are also a genetic by-product, caused by changes in gene(s) associated with other dental properties.

Genomic research has begun to explore genetic loci involved in enamel formation, mostly with the aim of understanding development, but also for inferences into genetic conditions (Paine et al., 2001; Hu & Yamakoshi, 2003; Hu et al., 2005; Ozdemir et al., 2005; Al-Hashimi et al., 2009). One such gene, ENAM, has been analyzed for differences among primate species (Kelley & Swanson, 2008). Evidence of adaptive ENAM evolution was implicated in observed interspecific variation in enamel thickness. Areas of this gene show signs of strong positive selection, probably related to enamel thickness changes (Horvath et al., 2014). Mutations in this same gene are responsible for many types of amelogenesis imperfecta (Crawford et al., 2007; Kelley & Swanson, 2008; Wang et al., 2015). Therefore, it is plausible that the genetic changes related to the evolution of extremely thick enamel and large posterior teeth in *Paranthropus,* over a relatively short period, created pleiotropy effects, including high rates of PEH.

In sum, comparisons of the frequencies and appearance of enamel hypoplasia defects among several fossil hominin and extant primate samples suggest that several potential causes for severe PEH on *P. robustus* molars can be ruled out. Given that: 1) substantially less defects are present on the species’ anterior teeth, 2) defects are relatively uniform in shape and size, and 3) all molars are similarly affected, the hypothesis that best fits the data is a genetic origin for ‘golf ball’-like PEH. A plausible scenario is that specific genetic mutations resulted in these defects and concerns the evolution of thick enamel and/or large posterior teeth in *Paranthropus*.

## Acknowledgements

The authors thank L. Berger and B. Zipfel from the University of the Witwatersrand, I. Livne from the Powell-Cotton Museum, and S. Potze from the Ditsong Museum of South Africa for access to their collections. For producing the CT scans provided by Ditsong Museum of South Africa we thank J.J. Hublin and the Department of Human Evolution, Max Planck Institute for Evolutionary Anthropology. For technical assistance we thank M. Skinner. This research was supported by a studentship to the first author from Liverpool John Moores University.

